# Single-cell multi-omics and spatial multi-omics data integration via dual-path graph attention auto-encoder

**DOI:** 10.1101/2024.06.03.597266

**Authors:** Tongxuan Lv, Yong Zhang, Junlin Liu, Qiang Kang, Lin Liu

## Abstract

Single-cell multi-omics data integration enables joint analysis of the resolution at single-cell level to provide comprehensive and accurate understanding of complex biological systems, while spatial multi-omics data integration is benefit to the exploration of cell spatial heterogeneity to facilitate more diversified downstream analyses. Existing methods are mainly designed for single-cell multi-omics data with little consideration on spatial information, and still have the room for performance improvement. A reliable multi-omics data integration method that can be applied to both single-cell and spatially resolved data is necessary and significant. We propose a single-cell multi-omics and spatial multi-omics data integration method based on dual-path graph attention auto-encoder (SSGATE). It can construct neighborhood graphs based on single-cell expression data and spatial information respectively, and perform self-supervised learning for data integration through the graph attention auto-encoders from two paths. SSGATE is applied to data integration of transcriptomics and proteomics, including single-cell and spatially resolved data of various tissues from different sequencing technologies. SSGATE shows better performance and stronger robustness than competitive methods and facilitates downstream analysis.

## 1. Introduction

Multi-omics typically includes the genomics, transcriptomics, proteomics, metabolomics, epigenomics, phenomics, and other single-omics [1, 2]. Research in these fields has enhanced our understanding of complex biological systems. Multi-omics data integration refers to feature representation and dimensionality reduction of different single-omics data to obtain their joint low-dimensional representation [3]. It can provide more comprehensive information for joint downstream analysis. One example is data integration of proteomics and metabolomics, which reveals novel connections between proteins and metabolites and successfully predicts the functions of five genes [4]. In addition, the data integration of transcriptomics and DNA methylation makes significant progress in biomedical classification tasks, offering new insights into the understanding and treatment of diseases [5].

The data integration of transcriptomics and proteomics is currently one of the focus point [6,7]. With the continuous development of sequencing technologies, it has become feasible to collect a large number of single-omics data. The emergence of spatially resolved transcriptomics and spatially resolved proteomics technologies provide access to the single-omics data with spatial information [8]. In particular, advanced sequencing technologies, such as SPOTS [9], spatial CITE-seq [10] and Stereo-CITE-seq [11], enable the simultaneous acquisition of transcriptome and proteome data on the same tissue section, thereby better maintaining the homology of different single-omics data. To utilize existing data in joint analysis and facilitate new discoveries in gene regulation, biological evolution, disease treatment, etc., the multi-omics data integration methods for transcriptomics and proteomics have become effective and important.

While multi-omics data integration plays a critical role in advancing research, it faces the challenge of addressing the inherent differences among different single-omics data. Sometimes these differences can be huge, requiring efforts on data consistency in terms of formats, dimensions, units, and normalizations [12]. Different single-omics data can be presented using various methods and tools to convey their biological significance. However, the integrated data still require reliable biological interpretation approaches [13].

Traditional data integration methods for transcriptomics and proteomics meet these challenges from statistics, network-based technologies and other perspectives. MOFA is a typical statistical method in an unsupervised fashion [14]. It infers a low-dimensional data representation in terms of interpretable factors that capture the global sources of variation across modalities, helping identify continuous molecular gradients and discrete sample subgroups. MOFA+ is an extension of MOFA [15]. It incorporates priors for flexible, structure regularization, enabling joint modeling of multiple groups and data modalities. Seurat v5, a network-based method, constructs a weighted nearest neighbor graph for data representation by connecting cells that share similarities across modalities, using learned cell-specific weights that determine the relative importance of different omics data [16]. Other traditional multi-omics data integration methods include SCIM [17], GRMEC-SC [18] and Mowgli [19]. Although these methods have made significant contributions, they are best suited for scenarios where the non-linear relationships between transcriptome and proteome data are not complex, and the challenges still remain.

With the continuous advancement of computing technology, the multi-omics data integration methods for transcriptomics and proteomics based on deep learning are emerging. TotalVI is a deep generative model utilizes a modeling strategy similar to scVI [20] to learn a joint probabilistic representation of paired gene expression and protein data [21]. ScMM addresses the complexity of multimodal single-cell data by using a mixture-of-experts multimodal variational autoencoder [22]. There are also other deep learning-based multi-omics data integration methods published, such as ScMDC [23], BABEL [24] and InClust+ [25]. These methods partially address the limitations of traditional methods, but there is still room for improvement in their performance. Additionally, almost all methods are designed for single-cell multi-omics data. When they are applied to spatial multi-omics data, fail to incorporate spatial information. The only method we have searched that involves spatial information is SpatialGlue [26]. However, it can not be applied on single-cell data, and its performance has not yet systematically verified.

Single-cell multi-omics data have the resolution at single-cell level that their integration enables comprehensive and accurate understanding of complex biological systems. Although spatial multi-omics data are usually more sparse, they provide spatial information that their integration facilitates the exploration of cellular spatial heterogeneity and more diversified downstream analysis. Therefore, it is necessary and significant to develop a reliable multi-omics data integration method that can be applied on both single-cell and spatially resolved data.

In addition to learning the representation of each single-omics data and obtaining an effective joint representation, multi-omics data integration also needs to retain the unique characteristics of each single-omics data. Graph attention auto-encoder (GATE) combines the graph attention mechanism with the auto-encoder [27], which has been verified to efficiently extract low-dimensional representations from complex graph-structured data [28]. This is crucial for mutil-omics data integration where initial data can be high-dimensional and sparse. The attention mechanism allows the model to selectively focus on important nodes and edges in the input graph, automatically learning which parts are critical for specific tasks [29], which is particularly advantageous in extracting important biological features from graph-structured data. As a self-supervised learning method, the auto-encoder framework is favorable for multi-omics data without ground truth labels [28]. Specifically, the graph neural network structure integrates information from neighboring nodes in the data through simple and often efficient calculations [30,31]. Notably, existing multi-omics data integration methods often focus on shared information among different single-omics data, neglecting specific information of each single-omics data. The dual-modality factor model can identify and extract both shared information across modalities and complementary information specific to each modality [32], which provides guidance for designing a model with dual-path framework to integrate multi-omics data.

This study presents a single-cell multi-omics and spatial multi-omics data integration method based on dual-path GATE (SSGATE). It can construct neighborhood graphs based on expression data and spatial information respectively, which is the key to its ability to process both single-cell and spatially resolved data. Two single-omics data are input into two GATEs via two separate paths. Two embeddings obtained through the encoders are integrated. Then they are used for reconstruction separately by the decoders. To train the model more effectively, a combined weighted loss of self-supervision and self-reconstruction losses [33] is adopted. SSGATE is applied on data integration of transcriptomics and proteomics, including both single-cell and spatially resolved data of various tissues from different sequencing technologies. Experimental results verify that SSGATE outperforms competitive methods in terms of performance and robustness. Additionally, SSGATE facilitates downstream analysis, such as cell clustering and trajectory inference.

## 2. Materials and methods

### 2.1 Datasets and preprocessing

The single-cell multi-omics datasets BMNC [34] and SLN111_D1 [21], and the spatial multi-omics dataset SCS_MT [11] are obtained (Table 1). They are converted to “h5ad” and “rds” formats for preprocessing and experiments. For transcriptome expression data, the count depth scaling with subsequent log plus one transformation is used for normalization, and then take the top 4000 highly variable genes to reduce the dimensions [35]. For proteome expression data, the centered log-ratio transformation is used for normalization [36].

**Table 1.** Details of multi-omics datasets (transcriptomics and proteomics).

| Type | Name | Tissue | Technology | Cell number | RNA number | Protein number |
| --- | --- | --- | --- | --- | --- | --- |
| Single-cell | BMNC | Human bone marrow | CITE-seq | 30,672 | 17,009 | 25 |
|  | SLN111_D1 | Mouse spleen | CITE-seq | 9,264 | 13,553 | 112 |
| Spatial | SCS_MT | Mouse thymus | Stereo-CITE-seq | 20,125 | 26,439 | 216 |

### 2.2 Dual-path graph attention auto-encoder for single-cell multi-omics and spatial multi-omics data integration

The overview of SSGATE is shown in Figure 1. Advanced sequencing technologies provide single-cell multi-omics and spatial multi-omics data. Neighborhood graphs are constructed for transcriptome and proteome data, respectively (Figure 1A). Two single-omics data are input into two separate GATEs for training through self-supervised learning, and finally, the integrated result is output for downstream analysis (Figure 1B).

**Figure 1.**
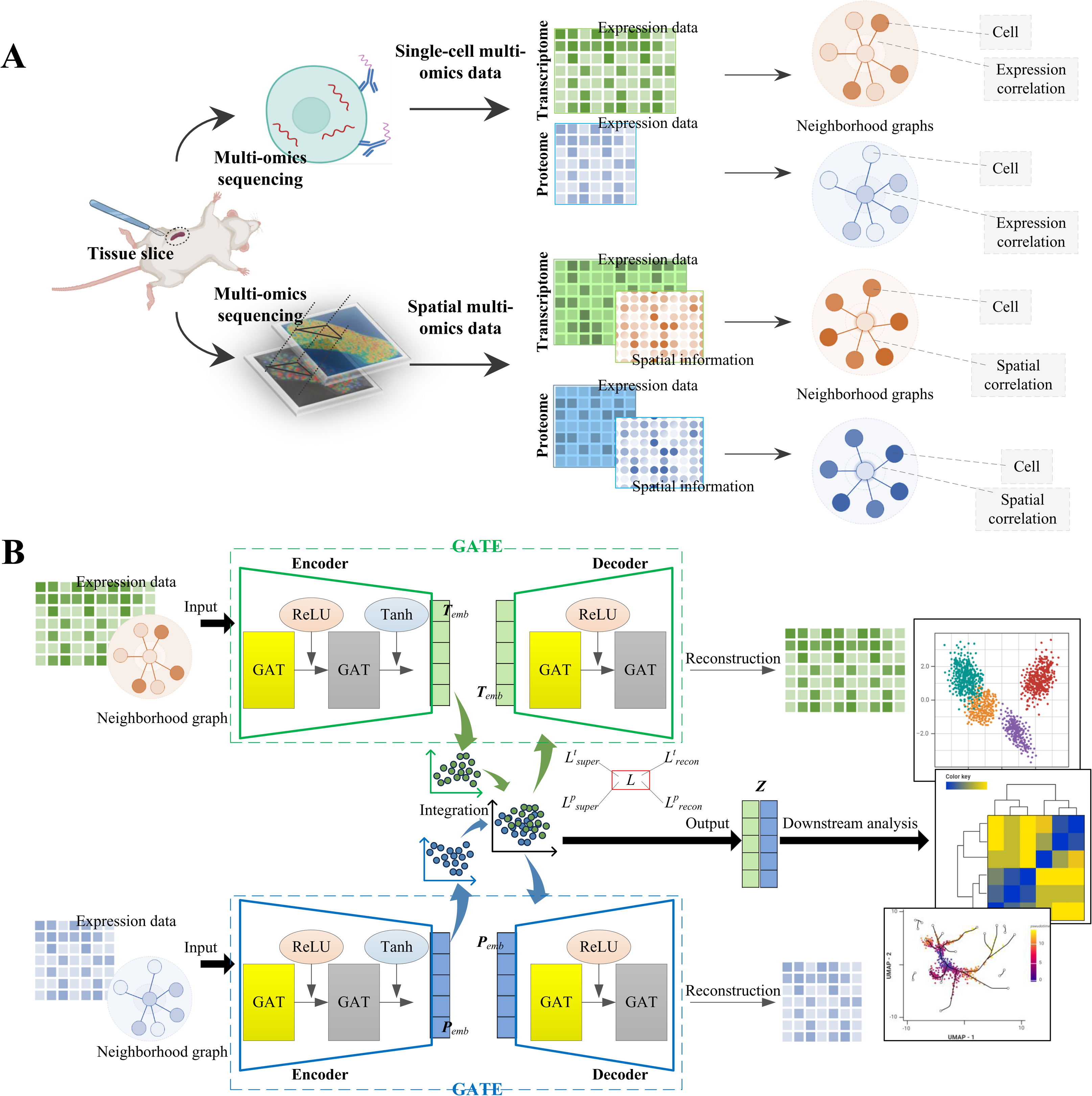
Overview of SSGATE. (**A**) Neighborhood graph construction for single-cell multi-omics and spatial multi-omics data. (**B**) Dual-path graph attention auto-encoder for multi-omics data integration. “GAT” represents the graph attention layer. “ReLU” and “Tanh” are the activation functions. ***T****_emb_* and ***P****_emb_* are the transcriptome embedding and proteome embedding, respectively. ***Z*** is the joint representation. *L* is the combined weighted loss. *L^t^_recon_* and *L^p^_recon_* are the transcriptome and proteome self-reconstruction losses, respectively. *L^t^_super_* and *L^p^_super_* are the transcriptome and proteome self-supervision losses, respectively.

#### 2.2.1 Neighborhood graph construction

For single-cell multi-omics data, the neighborhood graphs are constructed based on expression data, where each node represents a cell and each edge represents the expression correlation between two cells. The first step is the neighbor set generation. Due to the high dimensionality of transcriptome expression data, principal component analysis (PCA) [37] is used to reduce its dimensions to 200. For each cell *c_i_* (*i*=1,2,…,*m* and *m* is the total number of cells), its expression vector in transcriptomics is denoted as ***te****_i_*=[*te_i_*_,1_,*te_i_*_,2_,…,*te_i_*_,*t*_] and in proteomics as ***pe****_i_*=[*pe_i_*_,1_,*pe_i_*_,2_,…,*pe_i_*_,*p*_], where *t* and *p* are the total number of dimensions in transcriptomics and proteomics, respectively. The Euclidean distances between *c_i_* and *c_j_* are calculated as:

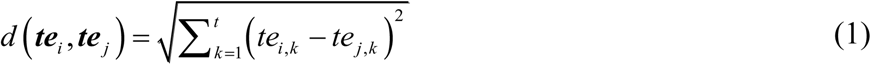

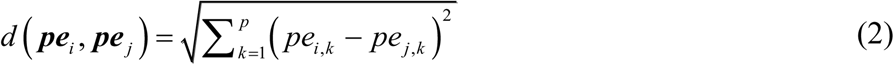

and the Euclidean distance sets of *c_i_* are obtained as:

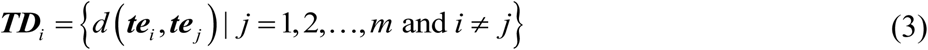

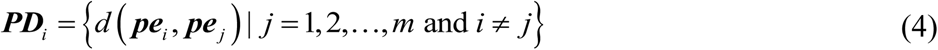

The neighbor sets of *c_i_* are generated as:

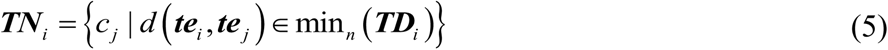

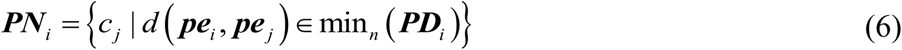

where *n* is the number of neighbors of *c_i_*. The second step is the neighbor pruning. All cells are clustered using the Leiden algorithm [38] based on transcriptome and proteome expression data, respectively. The neighbors of *c_i_* are pruned as:

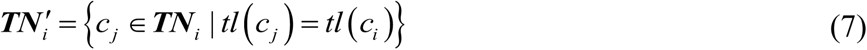

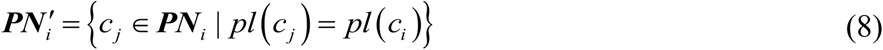

where *tl*(*c_i_*) and *pl*(*c_i_*) are the clustering labels of *c_i_* in transcriptomics and proteomics, respectively. The third step is the neighborhood graph construction based on neighbor sets of all cells as:

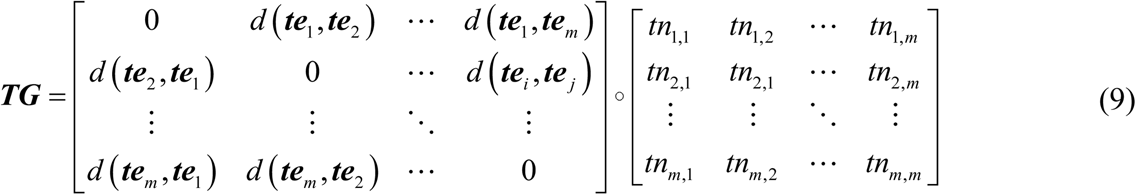

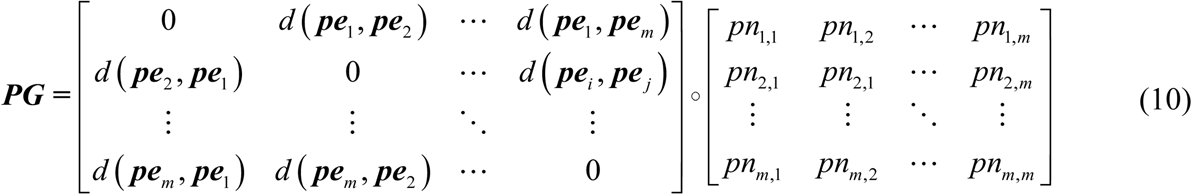

where ***TG*** and ***PG*** are neighborhood graphs in transcriptomics and proteomics, respectively, and

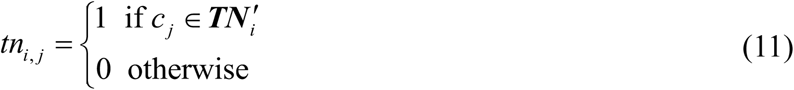

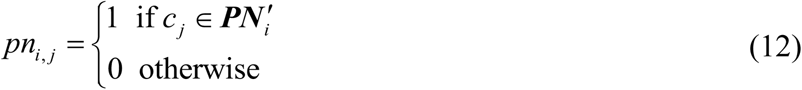

For spatial multi-omics data, the neighborhood graphs are constructed based on spatial information, where each node represents a cell and each edge represents the spatial correlation between two cells. The construction process is similar to it for single-cell multi-omics data, except that the coordinate data of the cells is used for calculating the Euclidean distances. The coordinate data contains more information on the relationships among cells and can better represent the distances among cells in real biological tissues.

#### 2.2.2 Dual-path graph attention auto-encoder architecture

The transcriptome expression data and neighborhood graph and proteome expression data and neighborhood graph are input into two separate GATEs. Each path’s GATE contains an encoder and a decoder. The encoder consists of two graph attention layers [28]. To maintain the capability to focus on important nodes and edges through the graph attention mechanism, while also preventing overfitting and conserving computational resources, the attention mechanism is activated in the first layer but deactivated in the second. The decoder adopts a symmetrical structure with the encoder. The “ReLU” and “Tanh” activation functions are used in encoders and decoders for nonlinear transformation.

In each epoch of training, each encoder encodes the inputs into a low-dimensional representation, i.e. embedding. The embeddings from the two paths are integrated as:

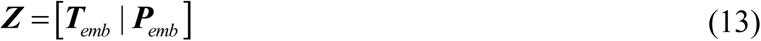

where ***T****_emb_* and ***P****_emb_* are the transcriptome embedding and proteome embedding respectively, and ***Z***, the joint representation, is used to calculate the self-supervision loss. Each decoder reconstructs the corresponding embedding into the original input to calculate the self-reconstruction loss. Self-supervision loss is the core for self-supervised learning, utilized for multi-omics data integration to preserve critical feature information from single-omics while ensuring that similar samples maintain their similarity in the joint representation. Self-reconstruction loss is employed to ensure accurate reconstruction of the original input from the joint representation, thereby preserving data integrity and enhancing model robustness. We adopt a combined weighted loss of self-supervision and self-reconstruction losses [33] to effectively train the model as:

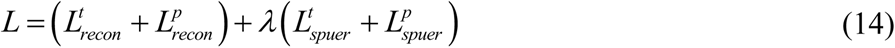

where *L* is the combined weighted loss, *L^t^_recon_* and *L^p^_recon_* are the transcriptome and proteome self-reconstruction losses respectively, *L^t^_super_* and *L^p^_super_* are the transcriptome and proteome self-supervision losses respectively, and *λ* is the balance parameter. The triplet loss function is used to calculate the self-supervision loss, and the mean squared error loss function is used to calculate the self-reconstruction loss. The model parameters are updated through back propagation. When the number of epochs reaches the preset maximum value, the training stops, and then the joint representation of multi-omics data, i.e. the integrated embeddings, is finally output.

### 2.3 Cell clustering and trajectory inference process

The transcriptome and proteome expression data are clustered using the Leiden algorithm [38] to calculate and examine the differentially-expressed genes (DGEs) and proteins, respectively. High confident DGEs are selected based on *p*-values of Wilcoxon test and “log fold change” values for Gene Ontology (GO) enrichment analysis [39], which reveals the primary functions of different cell types. Pseudo-time analysis of the transcriptome data is conducted to infer the developmental trajectories of cells. Expression genes of cells with different cluster labels are analyzed to identify gene modules. The spatial gene expression profiles for different modules are drawn, and the relative expression levels of gene modules in different cell types are quantified.

### 2.4 Evaluation criteria

For the single-cell multi-omics datasets, which provide the ground truth cluster labels of the cells, the integrated data is clustered using the Leiden algorithm [38] to obtain the cluster label of each cell, and then Purity (P), Homogeneity Score (HS), Adjusted Rand Index (ARI) and Normalized Mutual Information (NMI) are calculated to evaluate different methods [18,19]. To ensure reliable comparison and analysis, the “resolution” parameter of the Leiden algorithm is set to 0.1, 0.2, …, 1.0, allowing us to obtain statistical results from 10 sets of independent experiments. We also rank different methods in each metric to calculate Robustness Rank Aggregation (RRA) [40], allowing for a holistic assessment of their performances. For the spatial multi-omics dataset, which has no ground truth, we conduct a series of downstream analyses and visualize the analysis results to demonstrate the facilitative role of SSGATE.

### 2.5 Project implementation

All scripts are written in Python 3.8.14. Normalization of transcriptome data, extraction of highly variable genes, calculation and selection of DGEs and proteins, PCA, and clustering based on the Leiden algorithm are implemented through SCANPY (v1.9) [35]. Normalization of proteome data is achieved by replicating the normalization function of Seurat (v5) [16] using Python. GO enrichment analysis is performed using ClusterProfiler4 (v4.0) [41]. Pseudo-time analyses are conducted separately through Monocle 3 (v1.0.0) [42] and PAGA [43]. Gene module identification is carried out with Hotspot (v1.1.1) [44]. The hyperparameters for SSGATE are set as: training epochs at 300, embedding dimension at 30, balance parameter at 0.1, learning rate at 0.001, and number of neighbors at 15. The project is implemented on STOmics Cloud, utilizing the default computing resources under the “GPU CUDA” node.

## 3. Results and discussions

### 3.1 SSGATE’s performance is affected by the number of neighbors

The number of neighbors is a crucial hyperparameter in the neighborhood graph construction. We verify its effect on the performance of SSGATE to determine an appropriate value (Table 2).

**Table 2.** Effect of the number of neighbors on the performance of SSGATE.

| Dataset | $n$ | P | HS | ARI | NMI |
| --- | --- | --- | --- | --- | --- |
| BMNC | 10 | 0.9769 | 0.9372 | 0.4229 | 0.6746 |
|  | 15 | **0.9785** | **0.9432** | **0.4437** | **0.6865** |
|  | 20 | 0.9702 | 0.9323 | 0.4044 | 0.6665 |
| SLN111_D1 | 10 | 0.6637 | 0.5960 | 0.4079 | 0.6186 |
|  | 15 | **0.6742** | **0.6106** | **0.4319** | **0.6328** |
|  | 20 | 0.6715 | 0.6094 | 0.4083 | 0.6270 |

When the number of neighbors is set to 15, SSGATE achieves optimal results on all metrics on two datasets, particularly excelling in the ARI, where its advantages are most pronounced. The experimental results indicate that various neighborhood graphs can be constructed based on different numbers of neighbors, thereby affecting the performance of SSGATE. In subsequent experiments, the number of neighbors is set to 15 by default.

### 3.2 SSGATE outperforms competitive methods on single-cell multi-omics data integration

SSGATE is compared with four competitive single-cell multi-omics data integration methods, including MOFA+ [15], Seurat v5 [16], totalVI [21] and scMM [22], to verify its performance (Figure 2). TotalVI and scMM are deep learning based and widely used methods, and MOFA+ (with its latest release on 2024.04.09) and Seurat v5 represent state-of-the-art methods.

**Figure 2.**
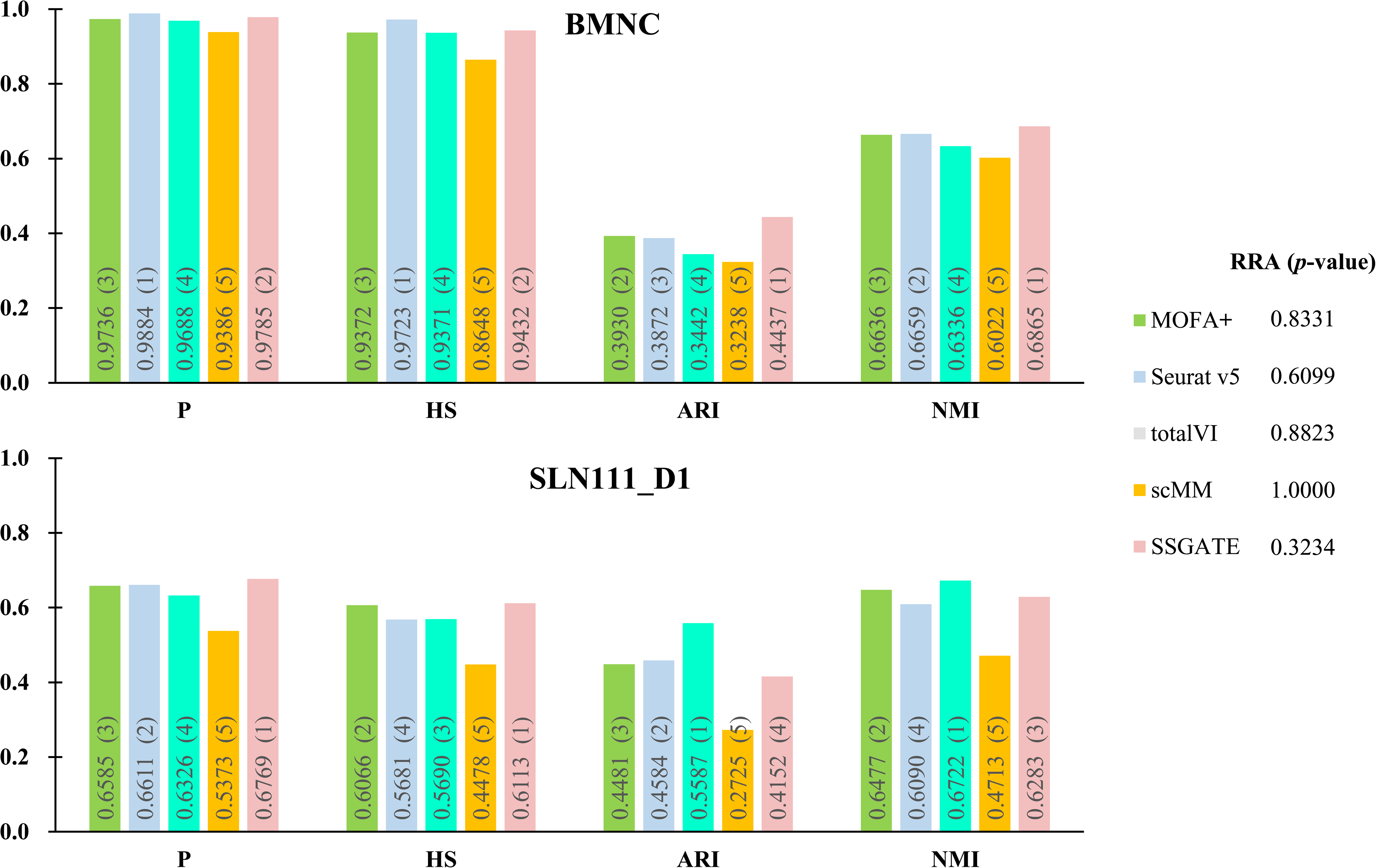
Comparison of SSGATE with other competitive methods on single-cell multi-omics datasets. Each result is the average value of 10 independent experiments. The numbers in parentheses represent the ranking of that value on the corresponding metric. The smaller the *p*-value of a method’s RRA, the more it indicates better overall performance.

On the BMNC dataset, scMM ranks fifth in each metric, with an average ranking of 5. Following is totalVI, which ranks fourth in every metric and has an average ranking of 4. MOFA+ ranks second in ARI and third in the other metrics, giving it an average ranking of 2.75. Seurat v5 ranks first in P and HS, second in NMI and third in ARI, resulting in an average ranking of 1.75. SSGATE ranks first in ARI and NMI and second in P and HS, with an average ranking of 1.5, making it the highest ranked method. On the SLN111_D1 dataset, scMM’s average ranking remains at 5. Seurat v5 ranks second in two metrics and fourth in two, with an average ranking of 3. MOFA+ ranks second in two metrics and third in two, and its average ranking is 2.5. TotalVI, ranks first in two metrics and ranks third or fourth in others, with an average ranking of 2.25. SSGATE ranks first in two metrics, ranks third in one and fourth in one, leading to the average ranking 2.25, tied with totalVI. Combining the rankings of all methods across all metrics on the two datasets, the *p*-values for RRA are calculated. ScMM has the highest *p*-value, followed by totalVI, then MOFA+, next is Seurat v5, and the smallest *p*-value is obtained by SSGATE. These results indicate that overall, SSGATE performs more strongly than other methods.

### 3.3 SSGATE shows strong robustness on single-cell multi-omics data integration with noises

SSGATE is also compared with the aforementioned four competitive methods on datasets with varying levels of Gaussian noise added [45] to verify its robustness (Figure 3).

**Figure 3.**
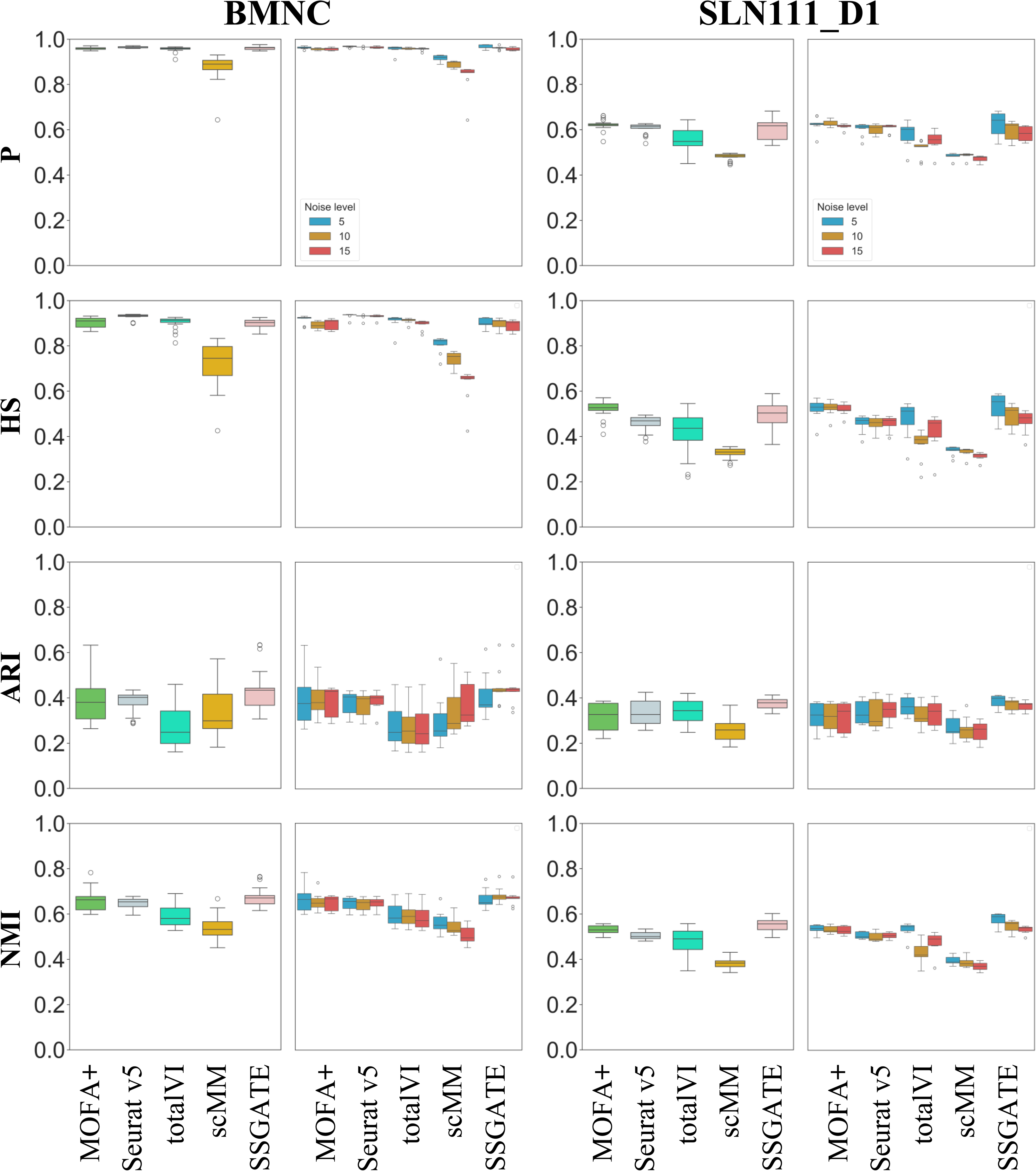
Comparison of SSGATE with other competitive methods on single-cell multi-omics datasets with varying levels of Gaussian noise added. On each dataset, the left column is the comprehensive results under three different levels of noise, and the right column is the separate results under three different levels of noise. Noise level 5, 10 and 15 represent 5%, 10% and 15% Gaussian noise added to the dataset, respectively.

From the separate results, the performances of all methods fluctuate when encountering different levels of noise, with SSGATE’s results being the best in most cases. Notably, scMM’s results exhibit more significant degradation with increasing noise than those of other methods in P and HS on the BMNC dataset, and totalVI’s results obviously fluctuate in P, HS and NMI on the SLN111_D1 dataset. Comprehensive results under three different levels of noise reveal that on the BMNC dataset, SSGATE achieves the best ARI and NMI, with its P and HS also close to the best results. On the SLN111_D1 dataset, SSGATE still obtains the best ARI and NMI. It also secures the second-best HS and the third-best P. Based on the comprehensive results, scMM, totalVI, Seurat, MOFA+ and SSGATE are ranked with average rankings of 4.88, 3.63, 2.38, 2.25, and 1.88, respectively, with SSGATE being the method with the highest average ranking. Although each method’s results exhibit the outliers, which may be due to insufficient stability causing some of the independent experiment results to deviate significantly from other results, SSGATE’s results contain the fewest outliers. For instance, in HS on the SLN111_D1 dataset, SSGATE is the only method with no outliers. These findings indicate that SSGATE can maintain excellent performance on the datasets with varying levels of noise, confirming its strong robustness.

### 3.4 SSGATE is applied to spatial multi-omics data integration, facilitating cell clustering and trajectory inference

The spatial multi-omics data from mouse thymus tissue is utilized to demonstrate the benefits of SSGATE for downstream analysis (Figure 4). From the UMAP plots (Figure 4A), SSGATE outperforms other methods in learning the integrated and discriminative latent space for tissue section, where cell clusters are more separated from each other. Additionally, the cell clusters from SSGATE’s results display strong spatial aggregation with clear boundaries, which is higher consistency with the fact that mouse thymus can be broadly divided into the outer region known as cortex and the inner medulla region [46]. We further demonstrate the effectiveness of SSGATE for combined DEGs and trajectory analysis. The GO-based enrichment analysis of DEGs reveals the progressive maturation, differentiation, and functional specialization of thymocytes within the mouse thymus (Figure 4B). Go terms associated with thymocyte differentiation and activation are prominently enriched in clusters 0, 1 and 5, such as T cell differentiation, B cell activation and activation of immune response. Clusters 2 and 4 exhibit GO terms indicative of mature lymphocytes, such as T cell mediated immunity and lymphocyte mediated immunity. In contrast, the significantly enriched Go terms in Cluster 3 are associated with supporting thymocytes development and maturation, such as anion and chloride transmembrane transports. These findings are in line with the fact that thymocytes start to develop in the outer region and migrate towards the inner region [47].

**Figure 4.**
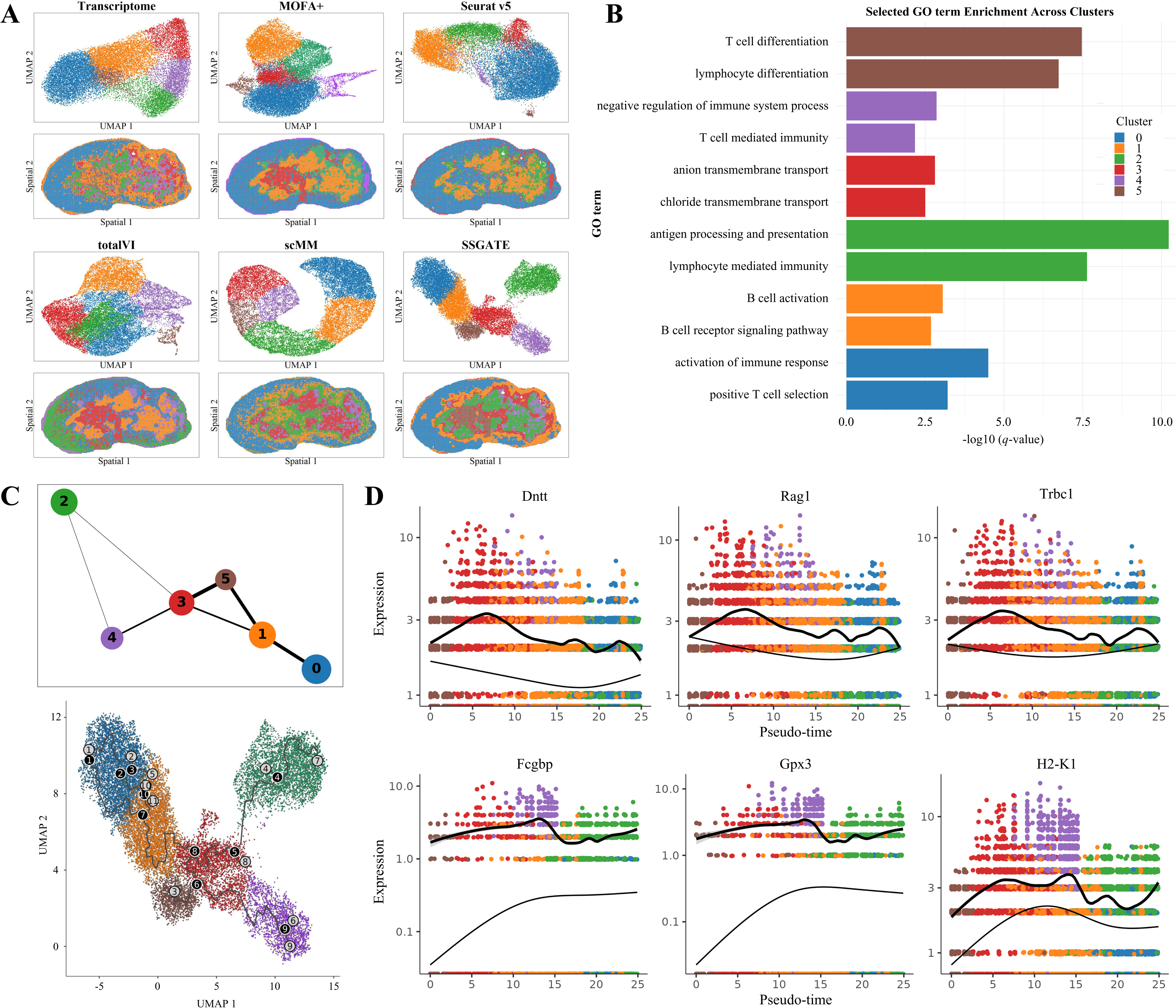
SSGATE is applied to spatial multi-omics data integration, facilitating cell clustering and trajectory inference. (**A**) UMAP plots for the integrated embeddings of SSGATE and other methods, and the spatial distribution of their identified cell clusters. (**B**) Top two highly enriched GO terms for top 100 ranked differently expressed genes of the identified cell clusters. (**C**) upper: PAGA graph of SSGATE embeddings, where each node represents a cell cluster that is connected by weighted edges that quantify the connectivity between clusters. Lower: Monocle 3 trajectory of the integrated cell clusters. (**D**) Pseudo-time kinetics of the significant genes varying along the inferred Monocle trajectory. For all subfigures, the cells are colored by the identified cell clusters, as in (**B**).

The PAGA graph of SSGATE embeddings exhibit a developmental trajectory from outer cortex to inner medulla region, showing high consistency with Monocle 3 results (Figure 4C). Furthermore, we identify genes that significantly vary over the inferred trajectory. In particular, the genes Dntt, Rag1, and Trbc1 are identified to be highly expressed during the early pseudo-time trajectory, and then exhibit a gradual decrease. In contrast, the genes Fcgbp, Gpx3, H2-K1 show an increasing trend and reached high expression levels in the later trajectory path (Figure 4D).

### 3.5 Discussions

SSGATE achieves the best overall performance primarily due to its ability to construct single-cell or spatial multi-omics data into complex graph-structured data. Through the GATE framework, it extracts low-dimensional representations and allows the model to selectively focus on important nodes and edges in the input graph. This is particularly beneficial for extracting important biological features. Additionally, the use of a combined weighted loss, although it may increase training time, effectively trains the model, enhancing its overall performance and stability.

As a self-supervised learning method, SSGATE can perform multi-omics data integration without requiring labeled data. It can adaptively recognize whether the input data is single-cell or spatial omics data and process it accordingly. Based on experimental experience, we provide a set of hyperparameters suitable for most datasets as default settings, minimizing the need for manual intervention, which facilitates usage by most researchers.

We also record the maximum memory usage of different methods in all experiments. High memory usage may result in method failures due to insufficient computational resources. The maximum memory usage of scMM is 1125 MB, the lowest among the methods, but its overall performance ranking is also the lowest. SSGATE’s maximum memory usage is 3351 MB. Next is totalVI, with a maximum memory usage of 5056 MB. Then comes MOFA+, with a maximum memory usage of 8259MB. Seurat has the highest memory usage at 12482 MB, which may because it typically requires separate normalization, dimensionality reduction, and other operations for each single-omics data before integration, involving a large amount of matrix computation. Considering both performance and computational resources, SSGATE is the most user-friendly method among these methods.

The integration of single-cell multi-omics data with high resolution facilitates more accurate cell classification and identification, while the integration of spatial multi-omics data with spatial information enables cell heterogeneity exploration and downstream analysis from a spatial perspective. Both types of multi-omics data have their characteristics and advantages, and as a method that can process both types of multi-omics data, SSGATE has significant potential for widespread applications.

## 4. Conclusion

This study proposes SSGATE, a single-cell multi-omics and spatial multi-omics data integration method based on dual-path graph attention auto-encoder. SSGATE constructs neighborhood graphs that effectively encapsulate expression data and spatial information, enabling it to process both single-cell multi-omics and spatial multi-omics data. The dual-path GATE model ensures that both shared and modality-specific information are meticulously preserved and utilized, enhancing the comprehensiveness of the data integration process. Experimental results demonstrate that SSGATE outperforms competitive methods in terms of performance and robustness and it facilitates downstream analysis. SSGATE provides researchers with a powerful tool to extract actionable insights from complex biological data. Future works will focus on optimizing the model’s workflow, improving its efficiency, and exploring its applicability to additional types of omics data.

## Code availability

The source code of SSGATE is available at https://github.com/Linliu-Bioinf/SSGATE.

## Acknowledgement

Project implementation was performed on the STOmics Cloud (https://cloud.stomics.tech).

